# Perturbed Mitochondrial Metabolism in Islets from Donors with Type-2 Diabetes

**DOI:** 10.1101/212548

**Authors:** Jiangming Sun, Ruchi Jain, Lotta E. Andersson, Anya Medina, Petter Storm, Peter Spégel, Hindrik Mulder

## Abstract

There is a preponderance for genes involved in ß-cell function among gene variants associated with future risk of type-2 diabetes (T2D). ß-cell function is controlled by metabolism of glucose, yielding signals triggering and amplifying insulin secretion. Perturbed ß-cell metabolism is a likely, albeit not proven, cause of T2D. We profiled metabolites in islets from T2D and non-diabetic donors, and found altered levels of mitochondrial metabolites in T2D. Analysis of genes encoding proteins localized to mitochondria (MitoCarta) by RNA-seq in an extended sample of islets revealed genes whose expression was associated with glycaemia- and/or BMI. Expression of two of these, α-methylacyl-CoA racemase *(AMACR)* and methylmalonyl-CoA mutase *(MUT),* was influenced by genetic variation (cis-eQTL). Silencing of *AMACR* and *MUT* in insulin-secreting cells reduced hormone secretion by 40-50%. In conclusion, by linking the metabolome to the transcriptome, we showed that perturbed mitochondrial metabolism is a feature of ß-cell dysfunction in T2D.

[Supplementary material is available for this article.]

## INTRODUCTION

Type 2 Diabetes (T2D) is a complex metabolic disease, involving perturbations in multiple metabolic pathways. Two primary disruptions are believed to underlie development of T2D: reduced insulin secretion from the pancreatic ß-cells and reduced insulin sensitivity in peripheral tissues (1). The relative contribution of these factors is debated (1). However, only ~20% of obese insulin-resistant individuals develop T2D (2). Thus, most obese subjects are capable of compensating for increased insulin resistance by enhancing secretion of the hormone from pancreatic ß-cells. This suggests that defects in the capacity to secrete insulin may be the ultimate triggers of the disease.

A concordance of T2D in monozygotic twins of 70%, compared with 20-30% in dizygotic twins, implies that T2D has a strong genetic basis (3). Genome-wide association studies (GWAS) of T2D and related metabolic traits have identified >100 variants associated with future risk of T2D (4). Among these, there is a preponderance for genes involved in ß-cell function (5, 6), favoring ß-cell failure as the ultimate cause of the disease.

Reduced ß-cell function and loss of ß-cell mass have been implicated in the impairment of ß-cell secretory capacity (7). Both these processes are controlled by cellular metabolism of glucose (8). When ß-cells are exposed to increased concentrations of glucose, the sugar is transported into the cell, metabolized in glycolysis and the Krebs cycle, leading to an increase in the cellular ATP/ADP-ratio. This rise in cellular energy charge triggers exocytosis of insulin (9). In addition, other coupling factors derived from cellular metabolism are thought to amplify secretion of the hormone (10, 11). Hence, T2D is likely to be associated with dysfunctional ß-cell metabolism (12). Indeed, abnormalities in mitochondrial morphology and function have been described in human islets from T2D donors (13). Moreover, we recently described a variant in the gene encoding transcription factor B1 mitochondrial *(TFB1M),* which is associated with reduced expression of mitochondrial encoded proteins, impaired insulin secretion and increased risk of future T2D (14, 15).

Given the implication of ß-cell failure in the pathogenesis of T2D and the importance of metabolism in regulation of insulin secretion, detailed investigations of metabolism in human islets promise to identify metabolic perturbations associated with ß-cell failure and development of T2D. Metabolomics is a powerful tool, enabling simultaneous analysis of levels of great numbers of metabolites (16). Integration of gene expression and metabolite profiling data, depicting the cellular machinery and its activity, respectively, may provide deepened understanding of cellular pathophysiology (17). In support of this approach, a recent study revealed that metabolic traits exhibit a stronger correlation to gene expression than to protein levels in mammals (18). The correlation between mRNA and protein levels, however, is generally poor (19), although models have been developed to regress mRNA levels with protein levels (20).

Here, we performed a detailed investigation of metabolism in human islets, using a top-down approach. Metabolites were profiled in a subset of islets from a large human islet gene expression data set. These analyses revealed perturbed mitochondrial metabolism in islets from T2D donors. The pin-pointing of mitochondrial metabolism prompted us to investigate expression of genes encoding proteins localized to the mitochondria, thus being likely to account for the changes in metabolism we observed. To this end, we used the MitoCarta gene inventory (21, 22), and identified genes that were targets of cis-expression quantitative trait loci (eQTL). The genes identified were associated with perturbed glycemia, both dependent on and independent of BMI, which was used as a marker for insulin resistance. To validate the functional significance of genes identified, two of these were silenced in INS-1 832/13 cells, resulting in reduced insulin secretion. Our results highlight the importance of mitochondrial metabolism in regulation of insulin secretion in general, but also imply a pathogenetic link between metabolic pertubations and expression of key metabolic genes in T2D.

## RESULTS

### Levels of mitochondrial-related metabolites are perturbed in islets from T2D donors

For metabolite profiling, we had access to islets from 70 donors, including 19 with T2D (Table 1). In these, we determined levels of 61 metabolites. First, we investigated differences in metabolite levels between islets from T2D and non-diabetic (ND) donors (Table 2 and Table S1). Lactate and alanine, end products of anaerobic metabolism, and Krebs cycle intermediates (succinate, malate and fumarate) were glucose-responsive in ND, but not in T2D, islets (Table 2). Levels of lactate and α-ketoglutarate, fumarate and malate were significantly elevated at basal glucose levels in islets from T2D donors (Table 2 and Figure 1). Hence, mitochondrial metabolism was clearly altered in islets from T2D donors: basal levels were elevated while glucose-elicited responses were lacking.

**Table 1:** Characteristics of islet donors. Date are expressed as average ± sd. Groups were compared using the two-tailed Student's t-test or Fisher exact test where appropriate.

|  | **Control (n=51)** | **Type 2 diabetes (n=19)** | **p-value** |
| --- | --- | --- | --- |
| Age (year) | 60.2±11.3 | 64.6±6.7 | 0.05 |
| Sex<sup>A</sup> | 19/32 | 8/11 | 0.79 |
| BMI | 27.0±3.9 | 27.6±4.6 | 0.64 |
| HbA1c | 5.8±0.5 | 6.8±1.0 | 0.0003 |
| Islet purity (%) | 68.2±18.7 | 70.8±23.6 | 0.66 |

<sup>A</sup> Sex is expressed as male/female.

**Table 2:** Glucose elicited alterations in metabolite levels in islets from donors with (T2D) or without T2D (ND). Only metabolites with significantly altered levels are shown. Additional information is found in Table S1.

| Catalog | Metabolite<sup>\$</sup> | ND<br>FC<sup>A</sup> | T2D<br>FC<sup>A</sup> | $P_g$ | $P_c$ | $P_x$ | $P_c$<br>(ND) | $P_c$<br>(T2D) | $P_g$ (LG) | $P_g$<br>(HG) |
| --- | --- | --- | --- | --- | --- | --- | --- | --- | --- | --- |
| Central cytosolic glucose metabolism | Gluc | 3.83 | 4.45 | 0.212 | <2.2e-16 | 0.177 |  |  |  |  |
| Central cytosolic glucose metabolism | FructP | 1.73 | 2.08 | 0.405 | 2.4E-08 | 0.159 |  |  |  |  |
| Central cytosolic glucose metabolism | Gluc6P | 1.08 | 1.46 | 0.182 | 0.006 | 0.238 |  |  |  |  |
| Central cytosolic glucose metabolism | SPAll | 1.03 | 1.17 | 0.134 | 2.8E-06 | 0.932 |  |  |  |  |
| Central cytosolic glucose metabolism | Lac | 1.02 | 0.82 | 0.123 | 1.9E-10 | 0.045 | 5.7E-09 | 0.283 | 0.0392 | 0.119 |
| Central cytosolic glucose metabolism | Ala | 1.11 | 1.03 | 0.232 | 3.6E-04 | 0.029 | 4.5E-04 | 0.763 | 0.0793 | 0.296 |
| TCA cycle | Cit | 1.02 | 0.93 | 0.677 | 0.002 | 0.506 |  |  |  |  |
| TCA cycle | IsoCit | 1.52 | 1.24 | 0.882 | 0.012 | 0.392 |  |  |  |  |
| TCA cycle | Succ | 1.15 | 0.99 | 0.434 | 0.007 | 0.023 | 0.001 | 0.643 | 0.11 | 0.748 |
| TCA cycle | Fum | 1.26 | 1.25 | 0.289 | 1.3E-05 | 0.0004 | 7.4E-07 | 0.273 | 0.036 | 0.945 |
| TCA cycle | Mal | 1.21 | 1.18 | 0.203 | 9.9E-07 | 0.022 | 5.1E-07 | 0.513 | 0.05 | 0.549 |
| Mitochondrial shuttles and shunts | Glu | 1.15 | 1.12 | 0.603 | 0.005 | 0.198 |  |  |  |  |
| Mitochondrial shuttles and shunts | Pglu | 1.25 | 1.05 | 0.608 | 0.035 | 0.070 |  |  |  |  |
| Mitochondrial shuttles and shunts | GABA | 1.31 | 1.07 | 0.471 | 0.003 | 0.684 |  |  |  |  |
| Amino acids | Gly | 1.08 | 0.98 | 0.720 | 0.341 | 0.032 | 0.065 | 0.019 | 0.566 | 0.600 |
| Amino acids | Ile | 1.08 | 1.02 | 0.594 | 0.460 | 0.011 | 0.519 | 0.059 | 0.184 | 0.969 |
| Amino acids | Lys | 0.96 | 1.04 | 0.335 | 0.007 | 0.026 | 2.9E-04 | 0.573 | 0.139 | 0.906 |
| Amino acids | Met | 1.05 | 1.00 | 0.877 | 0.276 | 0.028 | 0.842 | 0.026 | 0.504 | 0.368 |
| Amino acids | OHPro | 1.05 | 0.69 | 0.531 | 0.045 | 0.157 |  |  |  |  |
| Amino acids | Pro | 0.96 | 1.23 | 0.359 | 0.046 | 0.066 |  |  |  |  |
| Urea cycle | Orn | 1.09 | 1.07 | 0.896 | 0.005 | 0.187 |  |  |  |  |
| Lipid metabolism | C16:0 | 1.02 | 0.91 | 0.515 | 0.153 | 0.037 | 0.034 | 0.121 | 0.148 | 0.428 |
| Lipid metabolism | C20:0 | 1.17 | 1.11 | 0.978 | 0.963 | 0.039 | 0.360 | 0.580 | 0.398 | 0.260 |
| Miscellaneous | Fruct | 1.00 | 0.84 | 0.509 | <2.2e-16 | 0.898 |  |  |  |  |
| Miscellaneous | InosP | 0.98 | 0.88 | 0.257 | 0.267 | 0.046 | 0.085 | 0.379 | 0.154 | 0.865 |
| Miscellaneous | Creat | 0.97 | 0.83 | 0.905 | 0.026 | 0.209 |  |  |  |  |
| Miscellaneous | Udi | 1.07 | 0.95 | 0.695 | 0.017 | 0.436 |  |  |  |  |

**Figure 1.**
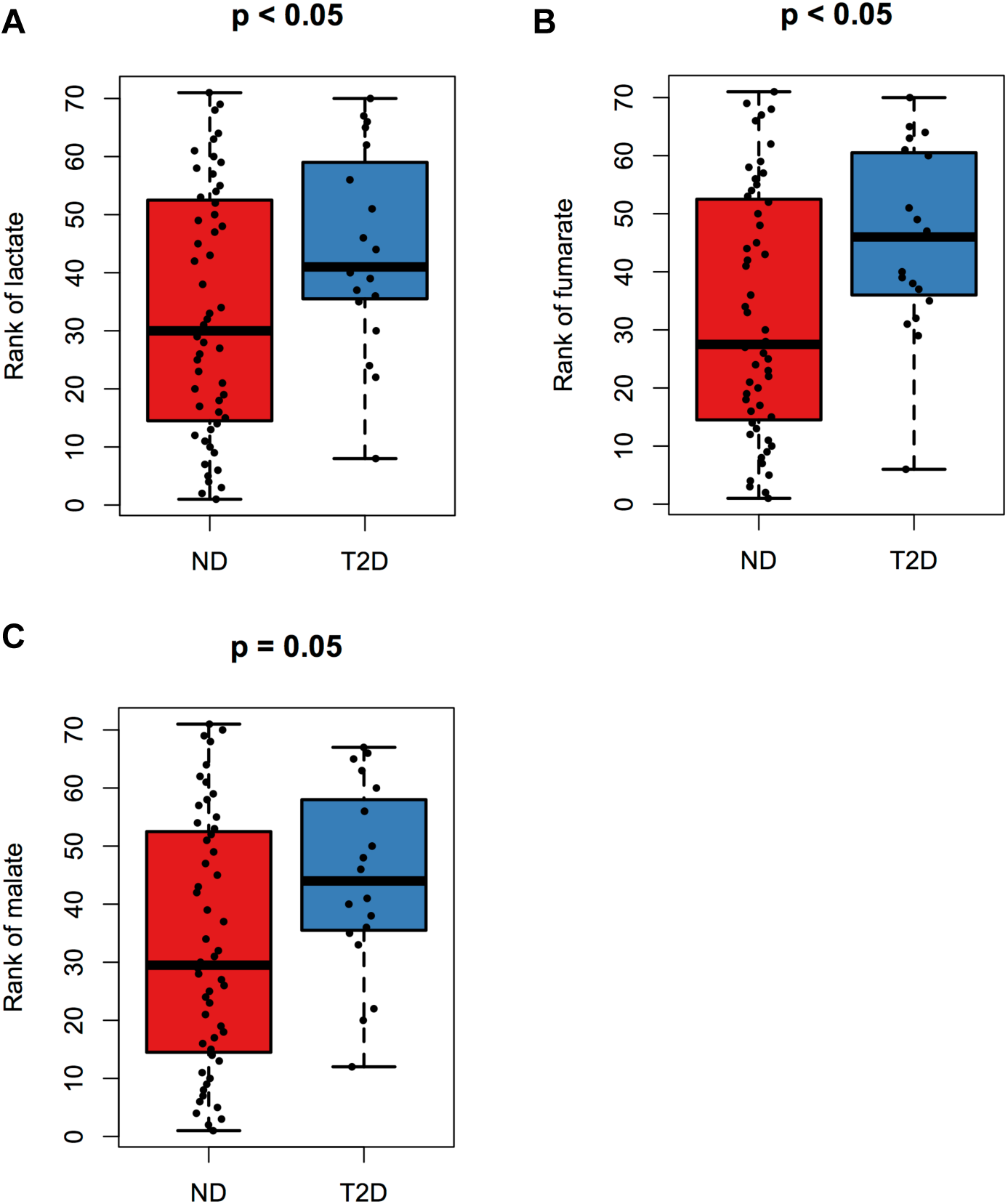
Alterations in basal metabolite levels. Basal levels of lactate and Krebs cycle intermediates fumarate and malate were elevated in islets from T2D donors. Islets from non diabetic (ND) donors (n=51) and T2D donors (n=19). P-values were obtained using an analysis of variance (ANOVA) adjusted for donor's age, sex and batch of experiments. Levels of metabolites were ranked in ascending order.

### HbA1c and BMI are associated with unique changes in expression of genes encoding proteins localized to the mitochondria

Gene expression in human islets has been extensively studied (23, 24), and expression data from 89 of the 131 islet batches examined here have previously been published (25). Given the perturbation in mitochondrial metabolism suggested by metabolite profiling, we focused our analysis on expression of genes encoding proteins involved in mitochondrial function. The MitoCarta inventory contains 1013 genes, which are homologues to 1098 mouse genes, encoding proteins that have been experimentally proven to localize to mitochondria (www.broadinstitute.org/pubs/MitoCarta) (21). After removal of genes expressed in less than 20 donors, 973 genes remained. Clinical data on the donors were limited due to ethical reasons (Table 1). This notwithstanding, in an attempt to tease out which pathogenetic process was responsible for changes in gene expression, we examined the associations between said genes and HbA1c and BMI; these were visualized in a two-dimensional plot (Figure 2). BMI was used as a surrogate measure for insulin resistance, whereas HbA1c reflected overall glucose tolerance, influenced by both insulin resistance and insulin secretion. Hence, this analysis allowed us to isolate changes in gene expression that were uniquely associated with either insulin resistance or insulin secretion: the assumption was that an association with HBA1c alone suggested impairment mainly of insulin secretion, an association with HBA1c and BMI suggested insulin resistance driving gene expression changes and, finally, association with BMI alone implied compensated normoglycemia. Thus, expression of 23 genes in human islets was found to be associated with both HbA1c and BMI, whereas expression of 81 genes were uniquely associated with HbA1c and 22 with BMI.

**Figure 2.**
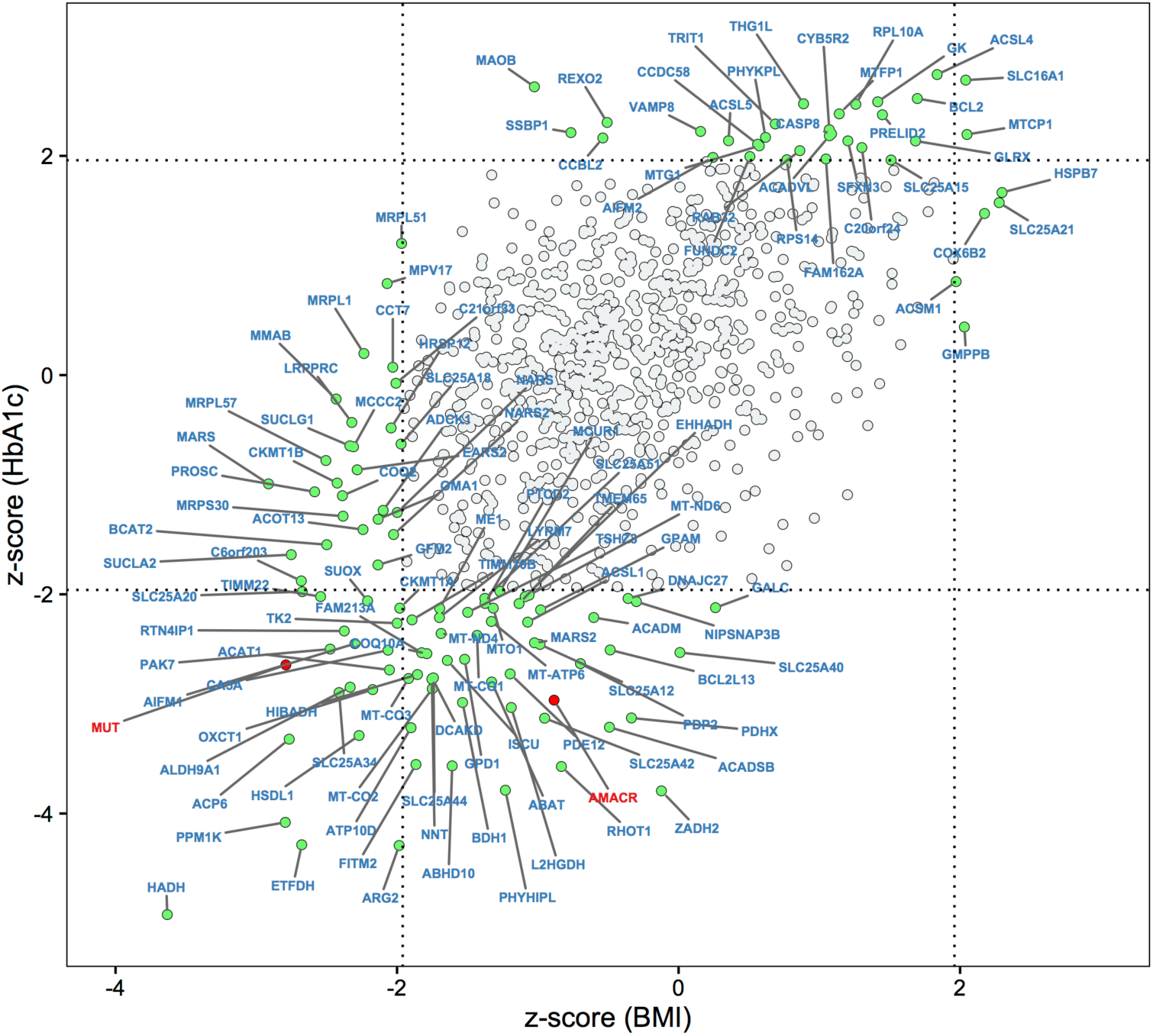
HbA1c and T2D differentially impact gene expression in human pancreatic islets. A plot of z-scores shows association between expression of genes encoding proteins localized to the mitochondria (MitoCarta) and HbA1c and BMI (n=131). Limits for significant associations (both absolute value of z-score>1.96 and permutation p<0.05) are indicated by dotted lines. Hence, genes with expression associated with both traits are found in the upper right and lower left quadrants. Genes uniquely associated with HbA1c are found in the center left and right quadrants and genes uniquely associated with BMI in the center top and bottom quadrants. Associations between gene expressions and phenotypes were examined, using ANCOVA, corrected by age and sex. The z-scores were converted from correlation coefficients by Fisher transformation. A permutation test (10000 permutations) was also performed by randomizing sample labels for gene expression data.

Metabolism of branched-chain amino acids (BCAA; KEGG: valine, leucine and isoleucine degradation) was enriched in the gene cluster that was inversely associated with HbA1c, BMI, and in both traits (Table 3). Moreover, expression of the monocarboxylate transporter 1 *(SLC16A1),* the latter being a ß-cell disallowed gene (26, 27), was positively associated with both HbA1c and BMI. Expression of NAD(P) transhydrogenase (*NNT*), which is deleted in a substrain of the glucose intolerant C57BL/6J mouse (28), was negatively associated with HbA1c. Associations of ribosomal proteins with HbA1c and BMI are shown in Table S2.

**Table 3:**
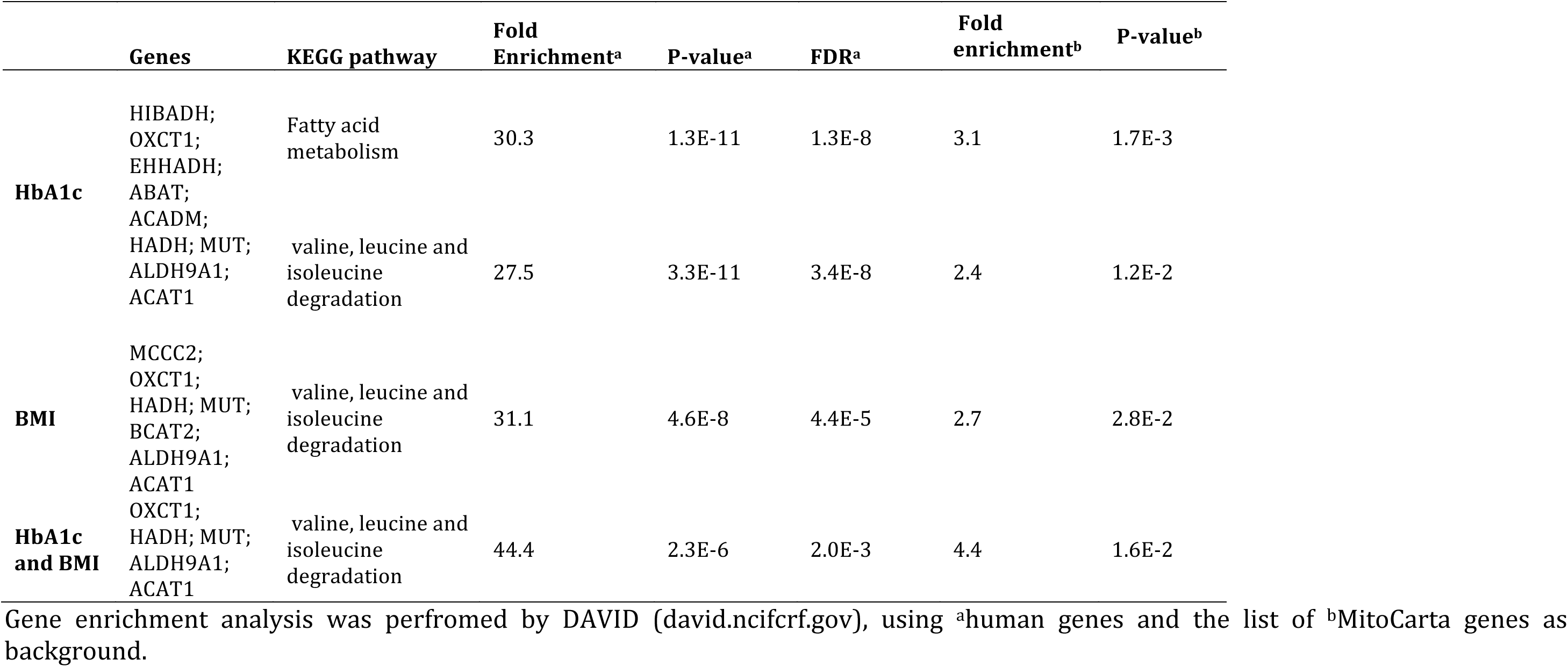
Pathway enrichment analysis of genes negatively associated with HbA1c, BMI and HbA1c/BMI.

### Expression quantitative trait loci (eQTLs)

Subsequently, we investigated whether expression of MitoCarta genes was impacted by genetic variations in islets from the 119 donors that had been genotyped. In total, we found 2458 cis-eQTLs, corresponding to 36 genes (Figure 3A and Table S3).

**Figure 3.**
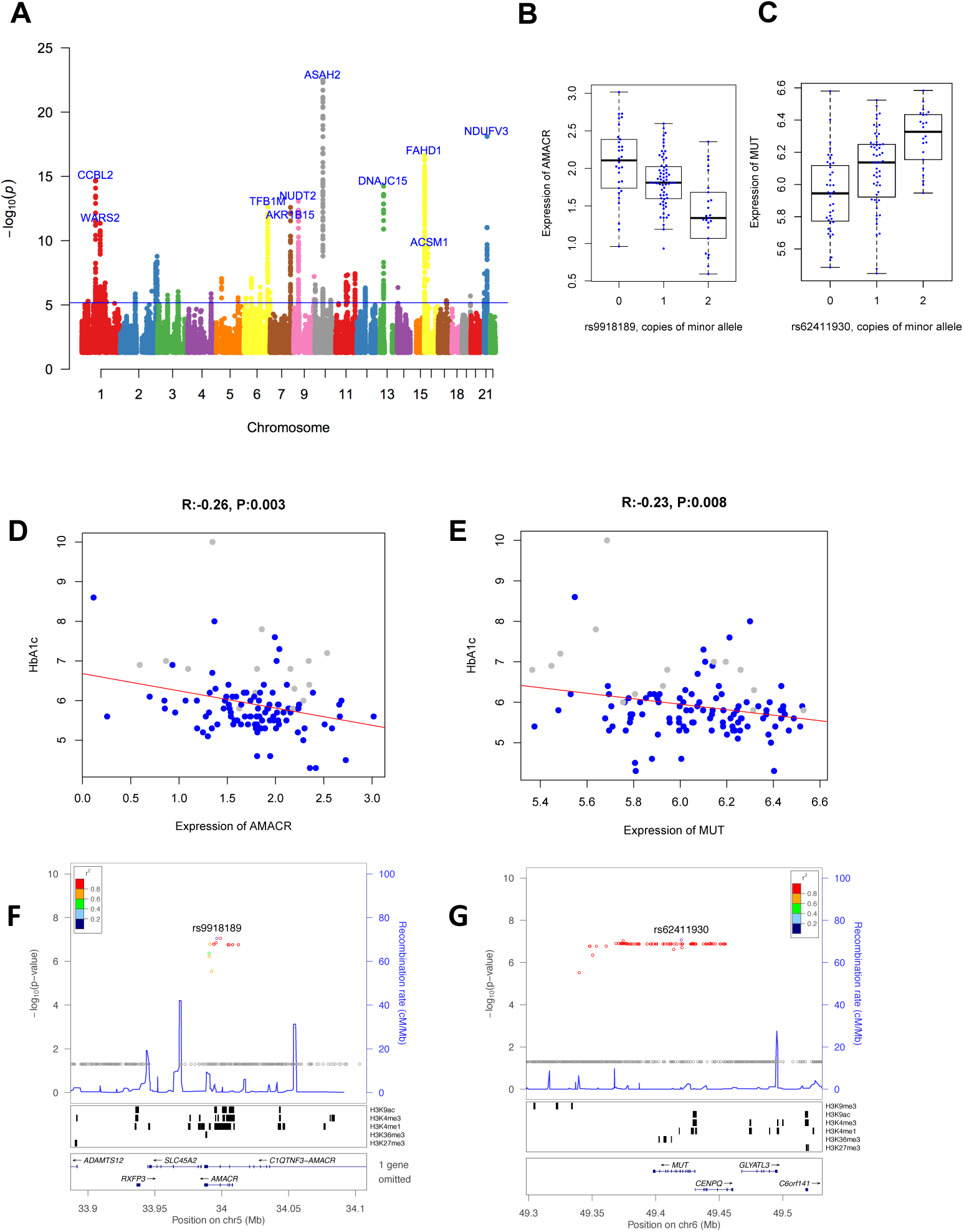
Impact of genetic variation on expression of genes encoding proteins localized to the mitochondria. **(A)** Manhattan plot of the best p-value per SNP, indicating the top 10 eQTL genes (n=119; permutation p<1.0e^−4^, F_1_-score>0.55). The blue line indicates FDR < 0.01**. (B-C)** Genotypes of eQTL SNP versus gene expression for *AMACR (p=8.9e*^*−8*^) and *MUT (p=8.5e*^*−8*^), respectively. Gene expression of *AMACR* and *MUT* are correlated with with HbA1c, adjusted by age and sex, with permuted p-value displayed in **(D)** and **(E)**, respectively, and their region plots shown in **(F)** and **(G)**, respectively.

Next, the eQTL SNPs were examined in the DIAGRAM (29) and MAGIC databases (30–32). The eQTL SNPs were enriched in T2D and glycemic traits, such as 2hr glucose, fasting glucose and HbA1c (Table S4). The eQTL genes were also examined in the T2D Knowledge Portal (T2D-GENES Consortium, GoT2D Consortium, DIAGRAM Consortium. 2017 June 1; http://www.type2diabetesgenetics.org/). We found seven genes *(PCCB, WARS2, TSFM, AGXT, SUCLG2, ACSM1* and *MRPL43)* showing genome-wide significant associations (p<5e^−8^) with height in the GIANT consortium (33). A locus-wide significant association (p<5e^−4^) with T2D was also found within the gene body of *MUT* in an European population (rs1141321, p=9.3e^−5^, effect size=0.7) (T2D-GENES Consortium, GoT2D Consortium, DIAGRAM Consortium. 2017 June 1; http://www.type2diabetesgenetics.org/gene/geneInfo/MUT). Islets with eQTL SNPs in *MUT* showed a strong trend for reduction in GSIS (p=0.051), and expression of *MUT* was lower in islets from T2D donors (nominal p=0.03).

Only seven eQTL genes (ACP6, *MUT, NARS2, ACSM1, AMACR, TRIT1* and *CCBL2)* showed phenotype-associated (BMI and/or HbA1c) expression (Figure S1), using our islet data. Of all eQTL SNPs within these genes, four non-synonymous coding mutations were found *(AMACR, MUT* and *NARS2;* Table S5). Moreover, a SNP in *AMACR* was predicted to be damaging to protein structure and/or function based on analysis by PolyPhen-2 (34). Lower expression of *AMACR* and *MUT* was associated with elevated HbA1c (Figure 3B-E).

The majority of eQTL SNPs were located in intronic regions (Table S6) and enriched in regions of islet active chromatin (Table S7). Notably, a large number of SNPs was found in the peak regions of histone modifications by H3K9ac and H3K4me3. As an example of this, the eQTL SNPs for *AMACR* and *MUT* were found in such regions (Figure 3F-G).

To confirm the validity of our results, we looked up eQTLs identified in islets in the Genotype-Tissue Expression (GTEx) pilot analysis (35). In total, 1475 cis-eQTLs corresponding to 27 genes *(ACP6, ACSM1, AGXT, AKR1B15, AMACR, ASAH2, CCBL2, DACT2, DNAJC15, EFHD1, FAHD1, FOXRED1, KIAA0141, MRPL21, MRPL39, MUT, NDUFA10, NDUFV3, NUDT2, PCCB, PITRM1, QRSL1, SARDH, TDRKH, TFB1M, TSFM and WARS2)* were reported as cis-eQTLs in GTEx (FDR<0.05) displaying the same direction of effects. A comparison of islet eQTLs and 44 tissue eQTLs is given in Table S8. Out of the shared cis-eQTLs, 63% (652 out of 1033) were found directionally consistent with GTEx data from human pancreas (Table S9). Seven out of 12 (58%) shared cis-eQTL signals were directionally consistent with a recent study (36).

### Expression of exocrine and endocrine cell markers and islet cell type-dependent gene expression

To ensure that islets from ND and T2D donors were comparable, we examined expression of exocrine and endocrine markers. We could not find any difference in the contribution from exocrine tissue between islets from T2D donors and ND donors, as judged by expression of the exocrine marker alpha 2 amylase *(AMY2A).* Moreover endocrine markers for specific cell-types, glucagon (*GCG*; α-cell), *MAFA* (ß-cell), and somatostatin (*SST*; delta cell) were similar between the groups. A nominally significant difference was observed in the ß-cell markers amylin *(IAPP;* p=0.04) and glucagon-like peptide 1 receptor *(GLP1R;* p=0.01) (Table S10).

Finally, we looked up HbA1c and/or BMI associated cis-eQTL genes identified in this work in seven recently published single cell studies (23, 37–42). Most of these cis-eQTL genes were not enriched in either α- or ß-cells, the dominating endocrine cell types, whereas conflicting results where obtained for some others. *SARDH* and *TDRKH* (38), *AGXT* (39), and *PITRM1* (41) have been suggested to be α-cell enriched in single studies. Two studies have revealed *EFHD1* to be mainly expressed in ß-cells (39, 42), whereas three studies have suggested *MAOB* to be ß-cell enriched (38, 39, 42). In two previous single cell studies, including islets from only six (38) and four (37) T2D donors, respectively, none of the genes identified in our study were found to be altered by T2D.

### Silencing of genes associated with lower HbA1c reduced insulin secretion

To examine whether the identified phenotype-associated cis-eQTL genes were functionally relevant, we performed gene silencing studies in a clonal insulin-producing cell line, INS-1 832/13 (43, 44). Among the phenotype-associated transcripts showing cis-eQTLs, we chose *AMACR* (Figure 3B and 3D) and *MUT* (Figure 3C and 3E) for further validation. These genes were negatively associated with HbA1c or HbA1c and BMI, respectively, but also found to be ubiquitously expressed in human islets. Moreover, *AMACR* and *MUT* have both been suggested to be expressed at moderate to high levels in pancreatic islets and clonal ß-cell lines at the mRNA (www.t1dbase.org), as well as protein, levels (www.proteinatlas.org). We attained a 78–86% reduction in *AMACR* mRNA levels, resulting in a pronounced 46-56% reduction in GSIS (p<0.01) (Figure 4). mRNA levels of *MUT* were similarly reduced by 70–82%, resulting in 31-36% reduction in secretion of insulin (p<0.01) (Figure 4).

**Figure 4.**
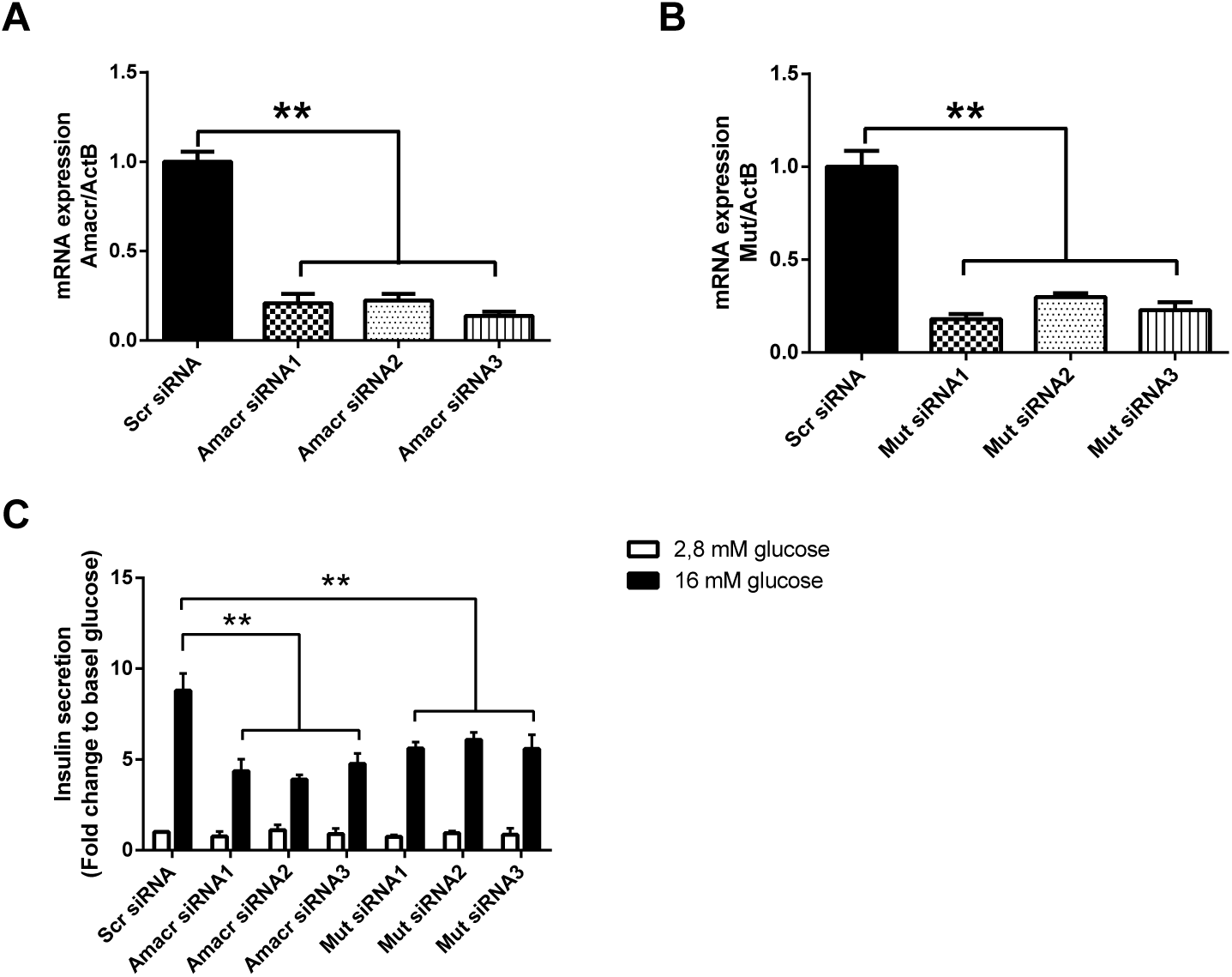
Silencing of the HbA1c-associated eQTL genes*AMACR* and *MUT* perturbs glucose-stimulated insulin secretion. Levels of *AMACR* **(A)** and *MUT* **(B)** mRNA were reduced by three different siRNAs per transcript. Knock-down efficiency was assesed by RT-PCR. **(C)** Basal insulin secretion (2.8 mM glucose) was unaffected whereas glucose-stimulated insulin secretion (16 mM) was significantly reduced after silencing of *AMACR* and *MUT.* Data are shown for three independent experiments for each siRNA and normalized to DNA content. Error bars represent mean ± SEM. **P < 0.01 versus control siRNA.

## DISCUSSION

Our study shows that mitochondrial metabolism is perturbed in islets in T2D: basal levels of metabolites were raised and glucose-elicited responses diminished. These findings highlight the importance of islet mitochondrial metabolism for proper ß-cell function and insulin secretion controlling whole body metabolism.

Given the pivotal role of mitochondrial metabolism, subsequent analysis of expression of genes encoding proteins localized to mitochondria revealed 75 and 38 transcripts negatively associated to HbA1c and BMI, respectively; 21 of these were shared. We assumed that genes associated with elevated HbA1c, but not BMI, reflected changes in insulin secretion. Changes associated with BMI, but not with HbA1c, reflected islet adaptation to an increased demand of insulin; when an association also existed with HBA1c, we assumed that islet adaptation to insulin resistance was insufficient, hence hyperglycemia. Thus, transcripts encoding proteins in BCAA metabolism were enriched among those associated with increased BMI. Indeed, elevated circulating levels of BCAA have been associated with obesity and insulin resistance (45), as well as an increased risk of future T2D (46).

Previous studies in human islets from patients with T2D showing mitochondrial dysfunction could not distuinguish between alterations being primary or secondary to the diabetic milieu (13). Altered mitochondrial transcriptional activity in response to elevated glucose and fatty acids, as well as oxidative stress, has been reported in ND islets (47). A strength of our study is that all islets were treated identically before analysis of glucose-elicited changes in metabolite levels. Responsiveness of cellular metabolism, and that of glucose in particular, to a rise in ambient glucose, is the fundament upon which stimulus-secretion coupling in ß-cells rests. For this reason, we believe that the impaired response in metabolite levels observed here are indeed part of the pathogenetic processes in T2D. Whether this is genetically or epigenetically driven remains to be determined. However, we have reported that a variant of the gene encoding TFB1M, a protein controlling mitochondrial protein synthesis, is associated with increased future risk of T2D mediated by ß-cell dysfunction. Together with the present findings, evidence for a primary role of mitochondria in T2D pathogenesis is mounting (14, 15).

Notable changes in transcription, besides those observed in BCAA metabolism, included higher expression of *MAOB* in hyperglycemic donors. Dopamine degradation, presumably catalyzed by MAOB, is increased in Parkinson's disease (48), a disorder associated with increased prevalence of T2D (49). Expression of *NNT* was uniquely decreased in islets from hyperglycemic donors, confirming previous findings in human islets (25). Out of the 1619 islet transcripts nominally associated with hyperglycemia (25), 63 encoded proteins localized to the mitochondria. Out of these, we verified 33 genes in the present study using 131 donors (52.4%).

Several eQTL SNPs were enriched in peak regions of histone modifications. This suggests that most variations will affect regulation of gene expression rather than protein structure. One of the eQTLs (rs950994) was linked to *TFB1M* expression (effect size: −0.34, FDR<0.005 and permutation p<1.0e^−4^), and has previosly been associated with reduced secretion of insulin in females (14). We observed four non-synonymous coding mutations in, *e.g., AMACR* and *MUT.* Out of the 36 cis-eQTL genes that we found, seven occur in genes with HbA1c- and/or BMI-associated expression.

Despite the importance of mitochondrial metabolism in GSIS, abrogation of mitochondrial genes does not necessarily result in disrupted hormone secretion (50, 51). Here, we found that silencing of either *MUT* or *AMACR* reduced GSIS. Both genes are expressed in multiple tissues, *MUT* being more ubiquitously expressed than *AMACR* (52) (www.proteinatlas.org). They are expressed at similar levels in adult α- and ß-cells, whereas *MUT* is more highly expressed in fetal α-cells, compared to ß-cells (42). MUT is a key enzyme in the conversion of the BCAAs isoleucine and leucine into the Krebs cycle substrate succinyl-CoA. Deficiency of MUT results in methylmalonic acidemia. Vitamin B-12 is a coenzyme of MUT. Notably, vitamin B12 deficiency, often associated with T2D (53), may cause methylmalonic aciduria (53, 54). Methylmalonate inhibits the Krebs cycle enzyme succinate dehydrogenase (55), which may account for its neurotoxic effects (56). Recently, expression of *MUT* in muscle was shown to be reduced in insulin resistance and T2D (57). This was associated with reduced levels of Krebs cycle intermediates, suggesting reduced Krebs cycle anaplerosis and flux, key processes in ß-cell stimulus-secretion coupling (58).

AMACR plays an essential role in degradation of branched-chain lipids, *e.g.,* phytanic acid (59). This process is initiated in peroxisomes, but AMACR is also required in mitochondria to manage remaining chiral methyl groups (60). *AMACR* has been associated with liver dysfunction (61) and cancer (62), the exact role of the corresponding protein in cancer being unknown. A mouse model of *AMACR* deficiency exhibits severely perturbed bile acid synthesis, but exhibited no clinical symptoms in absence of phytols in the diet (63). Phytanic acid, which is expected to accumulate upon AMACR deficiency, impairs mitochondrial energy-dependent activities (64).

There are some limitations in the study that need to be taken into account. Pharmacological treatment of T2D may have impacted islet function. Out of the 19 T2D donors, two had been treated with diet, whereas six were treated with metformin. Metformin reduces glucose production in the liver *in vivo,* where it is cleared by first pass metabolism. Indeed, most studies report no direct impact on insulin secretion (65, 66). Another limitation is that islets were obtained from diseased donors admitted to the intensive care unit. The average HbA1c of the ND donors is farily high (5.8%), which to some extent may depend on stress hyperglycemia (67) but also blood transfusions. A third limitation of our study is that we studied whole islets. It is therefore difficult to draw conclusions about specific cell populations. Moreover, proportions of, *e.g.,* α- and ß-cells, may vary. However, we did not find any difference in exocrine markers and only a very weak nominal difference in the ß-cell markers *IAPP* and *GLP1R.* Moreover, only two of the cis-eQTLs identified in our study, *EFHD1* and *MAOB,* have been replicated as ß- (39, 42) and α-cell enriched (38, 39, 42), respectively, in studies on sorted islet cells. This notwithstanding, MAOB has been shown to be expressed at the protein level in human ß-cells and to be involved in T2D-associated ß-cell dysfunction (68). Isolation, culture, dissociation of islets and sorting of islet cells differ between laboratories and may thereby affect cell-type specific gene expression (69). Moreover, islet dissociation will disrupt paracrine regulation in the islet (70). Hence, removing the ß-cell from its natural environment is also likely to impact on its function (71, 72). Finally, a potentially confounding factor in these analyses is reports on islet size to be increased in insulin-resistant individuals (73), and secretion of insulin from smaller islets being superior to that from larger (74).

We focused on alterations in metabolite levels in response to glucose stimulation, which should be less impacted by such differences. Importantly, purity did not differ between islets from ND and T2D donors in the present study. Islets were cultured for different times prior to RNA-Seq and metabolite profiling, precluding co-analysis of metabolite and gene expression data. Also, incubations performed prior to metabolite profiling were conducted in buffer, and not complex defined media, to improve detection of glucose-elicited changes in metabolism. Hence, to overcome these experimental differences, metabolite profiling and gene expression data were analyzed separately. To this end, our study, to our knowledge, represents the largest collection of human islets in which metabolism has been profiled, and showed that mitochondrial metabolism is pertubed in islets from T2D patients.

## METHODS

### Human islets

Pancreatic islets from deceased human donors were obtained from the Human Tissue Laboratory at Lund University Diabetes Centre, which receives islets from the Nordic Center for Clinical Islet Transplantation (http://www.nordicislets.com, Uppsala, Sweden). Islets were processed and their quality assessed as previously described in detail (25). In total, we received islets from 70 donors, including 19 with T2D, for metabolite profiling. The groups of ND and T2D individuals did not differ in age, sex or BMI; only HbA1c was higher in islets from T2D donors (6.8±1.0 in T2D and 5.8±0.5 in ND donors, p<0.001) (Table 1). In addition to this, we obtained RNA sequencing (RNA-Seq) data from 131 human islet preparations, out of which 70 were used for metabolite profiling and 119 were genotyped and used for detection of cis-eQTLs. The genome and transcriptome from 89 of these donors have previously been investigated (25).

### Metabolite profiling

Human islets were hand-picked under a stereo microscope and cultured overnight as described previously (25). Subsequently, 300 islets making up one sample were transferred to new dishes containing Hepes-balanced salt solution (HBSS) supplemented with 2.8 mmol/l glucose (75, 76). Islets were pre-incubated at 37°C in 95% air and 5% CO_2_ for 30 min, washed in PBS and further incubated for 1 hour in either 2.8 mmol/l glucose or 16.7 mmol/l glucose. Finally, islets were centrifuged (800xG, 4°C, 2 min), washed with 0.9% NaCl, centrifuged and snap froozen in liquid nitrogen. Islets were then stored at −80°C until analysis. Metabolites were extracted and analyzed on an Agilent 6890N gas chromatograph (Agilent Technologies, Atlanta, GA) connected to a LECO Pegasus III TOFMS electron impact mass spectrometer (LECO Corp., St. Joseph, MI), as previously described in detail (76, 77) (see also Supplemental Experimental Procedures).

### Gene expression

Gene expression was performed in islets from 131 donors using RNA-Seq (Illumina's TruSeq RNA Sample Preparation Kit and analyzed on the Illumina HiSeq2000 platform), as previously described in detail (25) (see also Supplemental Experimental Procedures).

### Expression quantitative trait loci (eQTL) analysis

Genotyping was performed on 119 human islets, using Illumina HumanOmniExpress 12v1 C chips. Methods for genotype calling, quality control and imputation have previously been described in detail (25) (see also Supplemental Experimental Procedures).

### Cell culture and Transfection

The clonal rat insulin-producing cell line, INS-1 832/13, was cultured in complete RPMI 1640 containing 11.1 mM glucose and supplemented with 10 % fetal bovine serum, 10 mM HEPES, 2 mM glutamine, 1 mM sodium pyruvate, and 50 μM ß-mercaptoethanol. The cells were maintained at 37 °C in a humidified atmosphere containing 95% air and 5% CO2. *AMACR* and *MUT* were knocked down using three different Silencer Select Pre-designed siRNA (Amacr siRNAl = s129370; Amacr siRNA2 = s129371; Amacr siRNA3 = s129372; Mut siRNAl = s153827; Mut siRNA2 = s153828; Mut siRNA3 = s191452; Ambion, USA) and lipofectamine RNAiMAX reagent (Invitrogen).

### RNA extraction and Quantitative real-time PCR

Total RNA was extracted from INS-1 832/13 cells, using RNeasy mini kit (Qiagen) according to the manufacturer’s protocol. One μg RNA was reverse transcribed using RevertAid First-Strand cDNA synthesis kit (Thermo Scientific) and quantitative real-time PCR was performed using the TaqMan gene expression assay (Amacr: Rn00563112 and Mut: Rn01512343; Applied Biosystems, Life Technologies), using a 7900HT Fast Real-Time System (Applied Biosystems). Gene expression was quantified by the comparative Ct method using ß-actin (Rn00667869) as reference.

### Insulin secretion assay

Insulin secretion from INS-1 832/13 cells was assessed by static incubation as previously described (78). Briefly, cells were preincubated in HEPES balanced salt solution (114 mM NaCl; 4.7 mM KCl; 1.2 mM KH_2_PO_4_; 1.16 mM MgSO_4_; 20 mM HEPES; 2.5 mM CaCl_2_; 25.5 mM NaHCO_3_; 0.2% bovine serum albumin, pH 7.2) supplemented with 2.8 mM glucose for 2 h at 37°C. Then cells were washed with PBS and incubated in the same buffer containing either 2.8 mM (low glucose, LG) or 16.7 mM (high glucose, HG) glucose for 1 h. Released insulin in the supernatant was assayed using an ELISA for human insulin (Mercodia) and normalized to total DNA using Quant-iT™ PicoGreen^®^ dsDNA Assay Kit (Thermo scientific), according to manufacturer’s protocol.

### Statistics

Glucose-elicited changes in metabolite levels were expressed as the ratio of their normalized peak area (A) at 16.7 and 2.8 mM glucose. Levels of metabolites were rank transformed prior to statistical analysis. The effect of glucose-elicited changes was determined by a linear mixed-effects model with the donor as a random effect while accounting for batch, age, sex, group (ND, T2D) and condition (LG, HG) as fixed effects. When a significant interaction between the condition and group was found, a linear mixed-effects model accounting for batch, age, sex and condition (LG, HG) as fixed effects was used to evaluate the effect of glucose-elicited change within ND and T2D islets, respectively; an analysis of covariance (batch, age and sex) was used to study differences between groups at LG and HG stimulation. A P value less than 0.05 was considered statistically significant. Analyses were performed using R.

### Study approval

The study was approved by the local ethics committee in Lund (Dnr. 2011263).

## AUTHOR CONTRIBUTIONS

ELA, RJ, AMB and PS conducted the experiments. JS analyzed data. PSt assisted in analysis of the data. PS and HM conceived the project and drafted the manuscript. All authors contributed in writing the final version of the manuscript.

## ACKNOWLEDGEMENTS

This study was supported by grants from the Swedish Research Council (14196-12-5), EFSD/MSD, the Novo Nordisk foundation, Swedish Diabetes Foundation, The Swedish Strategic Research Foundation, Crafoord, Hjelt, Lars Hierta’s Minne, Fredrik och Ingrid Thuring’s, O.E. and Edla Johansson’s Vetenskapliga, Åke Wibergs, Direktör Albert Påhlssons, and Magnus Bergvalls Foundations, and the Royal Physiographic Society. This study was also supported by equipment grants from the KAW Wallenberg Foundation (2009-0243), and funding from the European Union’s Horizon 2020 Research and Innovation programme under grant agreement No 667191

